# Sleep/wake changes in perturbational complexity in rats and mice

**DOI:** 10.1101/2022.08.02.502525

**Authors:** Matias Lorenzo Cavelli, Rong Mao, Graham Findlay, Kort Driessen, Tom Bugnon, Giulio Tononi, Chiara Cirelli

## Abstract

In humans, the level of consciousness can be assessed by quantifying the spatiotemporal complexity of cortical responses using the Perturbational Complexity Index (PCI) and related PCI^st^ (st, state transitions). These measures are consistently high in wake and rapid eye movement (REM) sleep and low in dreamless non-REM (NREM) sleep, deep slow wave anesthesia, and coma. The neuronal mechanisms underlying the reduction of PCI/PCI^st^ in unconscious states remain largely unexplored. The extent to which different cortical areas or layers contribute to these measures is also unknown. To address these questions, here we first validate the use of PCI^st^ in freely moving rats (8 males) and mice (12, 4 females) by showing that its values are lower in NREM sleep and slow wave anesthesia than in wake or REM sleep, as in humans. We then show that low PCI^st^ is associated with the occurrence of an OFF period of neuronal silence. Moreover, the stimulation of deep, but not superficial, cortical layers leads to reliable changes in PCI^st^ across sleep/wake and anesthesia. Finally, consistent changes in PCI^st^ can be measured independent of which single area is being stimulated or recorded, except for recordings in mouse prefrontal cortex. These experiments directly support the hypothesis that PCI^st^ is low when an OFF period disrupts causal interactions in cortical networks. Moreover, they demonstrate that, as in humans, PCI^st^ can be used for the reliable assessment of vigilance states in unresponsive animals, without the need to rely on behavioral outputs such as the righting reflex.

**Significance Statement:** The level of consciousness can be assessed in humans by measuring the spatiotemporal complexity of cortical responses using the Perturbational Complexity Index (PCI) and related PCI^st^. These measures discriminate between conscious and unconscious conditions with high sensitivity and specificity and work in unresponsive patients. However, the neuronal mechanisms underlying PCI/ PCI^st^ are largely unexplored. Moreover, since they reflect evoked responses from many cortical regions, it is unclear whether some areas or layers are more informative than others. Here we validate PCI^st^ in rodents, provide direct evidence for the underlying neuronal mechanisms, and show that reliable changes in PCI^st^ can almost always be obtained independent of which single area is stimulated or recorded, but only after stimulation of deep layers.

## Introduction

The Perturbational Complexity Index – PCI – was developed in humans as a tool to quantify the complexity of the cortical event related potentials (ERPs) triggered by transcranial magnetic stimulation (TMS) (Casali et al., 2013). The underlying rationale was provided by the integrated information theory of consciousness, according to which high complexity, resulting from the combined presence of high integration and high information in corticothalamic networks, is a prerequisite for being conscious (Tononi et al., 1994; Tononi, 2004; Tononi et al., 2016). In agreement with the theory, during wake, REM sleep, and ketamine anesthesia, when subjects retrospectively report vivid experiences, PCI is high because the stimulation triggers complex responses that are long-lasting and spread across many cortical regions. By contrast, during slow wave sleep and deep slow wave anesthesia, when subjects retrospectively report that they were not conscious, PCI is low because the response to stimulation either remains local, indicative of low integration, or is global but stereotyped, reflecting low differentiation (Casali et al., 2013). The analysis of PCI data from a benchmark population of 150 subjects in whom the presence or loss of consciousness could be established unequivocally led to the identification of an empirical PCI cutoff that, in healthy subjects, could discriminate between conscious and unconscious conditions with nearly 100% sensitivity and specificity (Casali et al., 2013). These results prompted the use of PCI to gauge the level of consciousness in patients in vegetative state (unresponsive wakefulness syndrome) and other difficult clinical situations where patients are either unresponsive or minimally responsive (Casarotto et al., 2016). Recently, another method to determine the complexity of the ERP was introduced, called PCI^st^, which calculates the overall number of non-redundant “state transitions” (st) caused by the stimulation (Comolatti et al., 2019). PCI^st^ has been validated using TMS and then extended to cases in which the cortical response was measured after deep intracranial electrical stimulation. It was found that PCI^st^ is nearly as accurate as PCI but easier and faster to compute, hence more suitable in clinical settings (Comolatti et al., 2019).

Although PCI and PCI^st^ are recognized as sensitive and specific measures to assess consciousness (Sarasso et al., 2021), the underlying cellular and network mechanisms are largely unexplored. In NREM sleep and slow wave anesthesia, when PCI/PCI^st^ values are low, cortical neurons are not tonically active but alternate more or less synchronously between ON periods of firing and OFF periods of silence (Steriade et al., 1993). It has been hypothesized that, under such conditions, cortical networks may be “bistable,” that is, they may not be able to support sustained causal interactions but, after brief periods of activity, necessarily fall into periods of silence. Under a bistable regime, strong stimuli are likely to silence the cortical network by triggering a large OFF period, thereby impairing causal interactions among cortical areas and resulting in a simple evoked response and low values of PCI/PCI^st^ (Pigorini et al., 2015; Sarasso et al., 2021). So far, however, studies in humans (Pigorini et al., 2015) and a recent study in anesthetized rats (Arena et al., 2021) could not provide direct evidence for the cessation of cortical unit firing underlying bistability.

In this study, we investigate PCI^st^ responses in freely-moving rodents—both rats and mice—using electrical and optogenetic stimulation accompanied by unit recording probes with multiple contacts (Neuropixels and NeuroNexus). We first demonstrate that PCI^st^ reveal changes in the complexity of neuronal responses with behavioral state—wake, NREM sleep, REM sleep, as well as anesthesia - that are similar to those observed in humans. Thus, PCI^st^ can serve as a reliable indicator of consciousness in laboratory animals, one that is highly validated in humans and that, unlike the standard righting reflex, can be employed in unresponsive states. We show that PCI^st^ changes can be recorded from a single site of penetration across multiple contacts, without the need to record from separate areas. We also show that the reduction of PCI^st.^ is associated with the triggering of neuronal OFF periods. We further investigate how different cortical areas and layers are involved in triggering and expressing complex neural responses.

## Materials and Methods

### Experimental animals

Adult rats (Sprague Dawley, males, 300–340 g; Charles River) and mice (CaMKIIα::ChR2 mice, both sexes, 19-28 g) were maintained on a 12 h light/12 h dark cycle with food and water available ad libitum (21–26 °C, 30–40% relative humidity). CaMKIIα::ChR2 mice were obtained by crossing CaMKIIα-Cre mice (Jackson Laboratory; T29-1; Stock No: 005359) with Cre-dependent ChR2(H134R)/EYFP expressing mice (Jackson Laboratory; Ai32; Stock No: 024109). All animal procedures and experimental protocols followed the National Institutes of Health Guide for the Care and Use of Laboratory Animals and were approved by the licensing committee. Animal facilities were reviewed and approved by the institutional animal care and use committee (IACUC) of the University of Wisconsin-Madison and were inspected and accredited by the association for assessment and accreditation of laboratory animal care (AAALAC).

### Surgical procedures

#### Rats

Stereotactic implant of the recording and stimulation electrodes was performed under isoflurane anesthesia (3% induction, 1.5-2.5% maintenance). Using sterile techniques, a midline incision was made to expose the skull and after cleaning the surface with bonding agent (OptiBond™), several small burr holes were made in the skull using a dental drill. Two stainless steel screws (0.8 mm tip diameter) were implanted to serve as ground for the stimulation probe (over contralateral olfactory bulb) and ground and reference for the recording probes (over the cerebellum). A 16-channel probe (NeuroNexus Technologies; A1×16-3mm-100-703-CM16LP) was implanted perpendicular to the cortical surface to be used as stimulation electrode, together with 1 or 2 Neuropixels 1.0 recording electrodes (Jun et al., 2017). The Neuropixels probes implanted in the frontal cortex (A/P +3.4, M/L +1.0 or A/P +4.2, M/L +2.0 angled towards the midline), reached secondary motor cortex (M2), prelimbic cortex (A32, PrL), or ventral/lateral orbital area (VO or LO). For simplicity, from here on we refer to VO and LO as orbitofrontal cortex (Of). In several animals, before insertion, the shank of the probes was coated with a red fluorescent cell-labeling solution (CM-Dil, Thermo Fisher Scientific) for later electrode track localization in postmortem histology. After probe alignment, insertion was performed using a robotic micromanipulator (New Scale Technologies) at a speed of 5 µm/s. At the end of the insertion the holes were sealed with silicone elastomer (Kwik-Sil™) and electrodes and probes were fixed to the skull using dental cement (C&B Metabond^®^). The implant was protected using a 3D-printed headcap based on the OpenEphys shuttleDrive enclosure (Voigts et al., 2019). At the end of surgery the margins of the implant were cleaned and antibiotic ointment was applied.

#### Mice

Surgery was performed under isoflurane anesthesia (2.0% induction; 0.8-1.5% maintenance) following sterile techniques. CaMKIIα::ChR2 mice of both sexes were implanted with optic fibers (Doric Lenses; core diameter = 200μm; NA = 0.22; diffuser layer tip) for optogenetic stimulations in either posterior (n = 9; 3 females) or anterior (n = 3; 1 female) cortex. For mice with posterior stimulation, optic fibers were placed on the cortical surface over the posterior parietal association cortex (PtA) (A/P -2.00, M/L ±1.80). For mice with anterior stimulation, optic fibers were implanted deeply in the cortex (A/P +1.93, M/L ±1.65, or A/P +1.77, M/L -0.60, angled towards the midline), to target anterior cingulate cortex (i.e., A24, Cg) and infralimbic cortex (i.e. A25, IL). For simplicity, from here on we refer to this targeted region together with PrL as mPFC (medial prefrontal cortex) (Carlen, 2017; Laubach et al., 2018; Le Merre et al., 2021).

To perform electrophysiology recordings, all mice were also implanted with laminar silicon probes (NeuroNexus Technologies; A1×16-3mm-50-177-CM16LP, A1×16-5mm-50-177-CM16LP, or A4×4-3mm-50-125-177-CM16LP), EEG and electromyogram (EMG) electrodes. To facilitate histological localization, in some cases the silicon probe shanks were coated with CM-DiI immediately before implantation. A right frontal silicon probe was implanted deeply in the cortex (A/P +1.93, M/L +0.40, or A/P +1.77, M/L +0.50, angled towards the midline), with electrodes targeting mPFC (Cg, IL or PrL). A left posterior parietal silicon probe was implanted in PtA (A/P -2.00, M/L -2.20), with electrodes targeting all layers. Reference screws were implanted over the cerebellum and olfactory bulb. EEG screw electrodes were implanted over left M2 (A/P +2.50, M/L -1.50) and right secondary somatosensory cortex (S2; A/P -1.30, M/L +4.0). EMG stainless steel wires were implanted bilaterally in the dorsal neck musculature and in the whisker musculature. The craniotomies and silicon probes were covered with surgical silicone adhesive (Kwik-Sil™), and all implants were fixed to the skull with dental cement (C&B-Metabond^®^, Fusio™ or Flow-It™ ALC™, Pentron).

### Experimental procedures and design

#### Rats

After surgery, all rats were kept in a temperature-controlled room (21-24 ° C) with a 12:12 light/dark cycle (light on at 9am) and with water and food available *ad libitum*. Rats were single housed in a transparent recording box (53 × 32 × 46 cm) containing bedding material and fully enclosed in a Faraday cage. After at least one week of recovery, the probes were connected to the recording system through a protection spring linked to a commutator, to allow free movements. After two days of adaptation, continuous recordings were performed for at least 48 hours to evaluate the sleep/wake pattern in baseline conditions, followed by 6 hours of sleep deprivation with novel objects (9am to 3pm) and subsequent sleep rebound (3pm to 9am the next day) to assess the homeostatic response to sleep loss (see OFF period analysis). Several days after sleep deprivation the stimulation sessions started and were conducted only during the light period, between 10am and 8pm, during wake, NREM sleep, REM sleep, as well as during anesthesia with sevoflurane (2%) or dexmedetomidine (0.1 mg/kg ip). Most of the rats were exposed to both anesthetics in a randomized order. Electrical stimulation of the cortical tissue was performed by delivering a vertical, 200-300 μm, bipolar, monophasic, current pulse of 0.5 ms of various intensities (30-100 μA). The depth of stimulation varied across experiments, always with cathode ventral. In each animal, the initial stimulation amplitude was set as the weakest one capable of triggering a slow wave during NREM sleep (Vyazovskiy et al., 2009a). Local field potentials (LFPs) and behavior were continuously monitored by the experimenter and stimuli were delivered only during consolidated episodes of wake and sleep, with the final goal of collecting 110 pulses for each behavioral state. In each session, stimuli were spaced apart at least 10 seconds, with often longer intervals to avoid sleep/wake state transitions. Shorter intervals (at least 4 secs) were sometimes used for REM sleep, whose bouts normally last approximately100 secs and account for less than 10% of the total behavioral time (Gonzalez et al., 2018). In each rat stimulation sessions spanned 2 to 4 weeks, interleaved with resting periods of at least 48 hours.

#### Mice

After surgery, all mice were kept in a temperature-controlled room (24-26 ° C) with a 12:12 light/dark cycle (light on at 10am) and with water and food available *ad libitum*. Mice were individually housed in transparent plastic cages (Allentown Caging; 24.5 × 21.5 × 21cm). The implanted silicon probes, electrodes and optic fibers were connected to the recording/stimulation system around one week after surgery to allow for recovery. Baseline recordings were acquired after the mice were accustomed to the system, then the experiments started after the temporal organization of sleep and wakefulness had normalized. All stimulation sessions were conducted during the light period. The implanted optic fibers were connected to a blue laser station (473nm, OEM Laser Systems DPSSL Driver, 100mW), which is triggered by the TDT system, with the laser output power manually controlled by an analog control knob on the driver. Based on the excitation threshold of specific opsins (Nagel et al., 2005; Madisen et al., 2012; Sidor et al., 2015), and the intended activation radius in the target area, the laser power to start with was estimated based on an established online calculator (https://web.stanford.edu/group/dlab/cgi-bin/graph/chart.php), which is modeled based on direct measurements in mammalian brain tissue. Laser power ranged from 0.2 to 2.9 mW at the tip of the optic fiber, and laser trains (8ms pulse width, 2000ms off between pulses, around 15 pulses per train) were delivered. Like in rats, the initial stimulation amplitude in each mouse was set as the weakest one capable of triggering a slow wave during NREM sleep. Behavior and electrophysiology data were continuously monitored by the experimenter, and stimuli were delivered during wake, NREM sleep, REM sleep, and under anesthesia with sevoflurane (1.0-2.0%) co-administered with dexmedetomidine (70-100 μg/kg, IP). The final goal was to collect around 80 pulses for each consolidated behavioral state.

### Histology

#### Rats

At the end of the last recording session, under general anesthesia (isoflurane 2-3%), rats were intracardially perfused with PBS (phosphate buffer solution with heparin 5000 IU/l) and 4% paraformaldehyde (PFA) in PBS for tissue fixation. Brains were then extracted and processed for histology. After fixation, brains were cryoprotected by exposure to increasing concentration of sucrose in PBS solutions at 4°C. Brains were then quickly frozen and sliced in coronal sections (40-50 μm thick) with a cryostat (Thermo Fisher Scientific; CryoStar™ NX50). Sections were dried overnight and mounted with medium containing DAPI (SouthernBiotech™; DAPI-Fluoromount-G). In some animals in which CM-DiI was not applied the sections were subjected to cresyl-violet (Nissl) staining. To verify the probe location sections were imaged with an upright epifluorescence microscope (Leica; DM2500).

#### Mice

To verify opsin expression, recording and cannula locations, mice were transcardially perfused under deep anesthesia (3.0% isoflurane, with a flush (∼30 s) of saline followed by 4% paraformaldehyde (PFA) in phosphate buffer (PB). Brains were removed and postfixed for 24 h in the same fixative, then cut in 50um thick coronal sections on a cryostat (CryoStar™ NX50 or Leica CM1900) after cryoprotection and flash-freezing. Sections were collected in PBS, mounted, air-dried, cover slipped (DAPI-Fluoromount-G, Vectashield, or Permount) and examined under a fluorescent or confocal microscope (Leica, Olympus). In some animals, to localize the silicon probes without fluorescent dye coating, glial fibrillary acidic protein (GFAP) staining was performed (rabbit-anti-GFAP primary antibody, DAKO Z0334, 1:1000 in blocking solution; Donkey-anti-Rabbit AF594 secondary antibody, 1:500 in blocking solution). In some cases Crystal Violet staining was performed to better visualize the location of the cannulas. To characterize the opsin expression of the CaMKIIα::ChR2 mice, in pilot experiments in 2 mice eYFP amplification staining was performed (rabbit anti-GFP primary antibody, Invitrogen, A11122, 1:1000 in blocking solution; goat anti-rabbit Alexa-488 conjugated secondary antibody, Invitrogen, A11008, 1:1000 in blocking solution).

### Electrophysiological recordings and stimulation

#### Rats

Electrophysiological recordings were performed using available Neuropixels 1.0 acquisition hardware (Putzeys et al., 2019). Neuropixels probes consist of a single shank (70 µm wide; 24 µm thick, 10 mm long) with 960 electrodes (2 columns; inter-electrode distance 20 µm), of which 384 can be recorded simultaneously (neuropixels.org). All experiments used the same electrode mapping, with a simple column expanding for 7.64 mm starting at the tip of the probe. Probes were connected to a head stage that transmit the data to a PXIe acquisition module mounted in a PXI chassis (National Instruments; PXIe-1071 chassis). The SpikeGLX software was used to acquire and visualize the data (https://github.com/billkarsh/SpikeGLX). In each probe the signal was amplified (x 500), digitized (10 bits) and filtered in two bands, one for the LFPs (0.5-500 Hz) and one for action potentials (AP; 0.3-10 kHz). LFP and AP signals were digitized at 2.5 and 30 kHz respectively with some small variation applicable after in brain calibration. Electrical stimulation was performed using a battery- powered 32 channels microstimulator system (Tucker-Davis Technologies; IZ2-32) connected to the 16 channels probe throughout a passive head stage and controlled with an electrophysiological recording software (Tucker-Davis Technologies; Synapse). Stimulus parameters and applied currents were recorded simultaneously in all channels.

All sessions were recorded with video (White Matter LLC; e3Vision system), and all data streams (video, stimulation, electrophysiology, etc.) were synchronized off-line using digital barcodes as described by the DAQ Synchronization Project from the Optogenetics and Neural Engineering Core at the University of Colorado Denver (https://optogeneticsandneuralengineeringcore.gitlab.io/ONECoreSite/projects/DAQSyncronization/).

#### Mice

Electrophysiological recording and optogenetic stimulation were performed using RZ2 BioAmp processor and OpenEx software (Tucker-Davis Technologies). Silicon probes were connected through a head stage to an amplifier (Tucker-Davis Technologies; PZ5 NeuroDigitizer Amplifier) before reaching the RZ2 processor. EEGs and LFPs were filtered by 0.1-100Hz, and multi-unit activities (MUAs) were filtered by 0.3-5kHz. Sampling rate for storage was 256Hz for LFPs, EEGs and EMGs; 25kHz for MUAs. Spike data were collected discreetly from the same LFPs channels.Amplitude thresholds for online spike detection were set manually based on visual control. Whenever the recorded voltage exceeded a predefined threshold, a segment of 46 samples (0.48 ms before, 1.36 ms after the threshold crossing) was extracted and stored for later use. All sessions were recorded with video.

### Sleep scoring and data processing

Sleep scoring was performed manually using a fork (https://github.com/TomBugnon/visbrain) of Visbrain Sleep (Combrisson et al., 2019), which includes several enhancements to facilitate the scoring of non-human sleep. Analysis of electrophysiological data was performed in MATLAB R2019b (The MathWorks^®^). LFP data were visually inspected to remove artefacts. Isolated bad channels were replaced by the mean of the immediately surrounding good channels. All LFP channels were subjected to linear detrend and lowpass filtering (200 Hz), using a zero-phase distortion third order Butterworth filter. Single trials were extracted in a ±4 sec window using stimulation time as zero. All trials and channels were visually inspected (SpikeGLX). Trials were discarded if there were artifacts in the few seconds around the stimulus, or when the stimulus was delivered close to a sleep/wake transition.

### PCI^st^

The spatiotemporal complexity of cortical event related potentials (ERPs) was quantified using a variant of the original perturbational complexity index or PCI (Casali et al., 2013; Sarasso et al., 2015), called the PCI state transition (PCI^st^) variant (Comolatti et al., 2019). With this method the principal components accounting for at least 99% of the variance present in the ERP response are obtained through singular value decomposition and then selected based on their own baseline level (signal-to-noise ratio, SNR_min_). The number of state transitions (NST) is then measured for each principal component during baseline (-800 to -100 msecs) and after the stimulus (10 to 800 msecs). NST is a measurement adapted from recurrent quantification analysis over the ERP distance matrices (Comolatti et al., 2019). PCI^st^ is the sum of the differences in NST between the baseline and the response for each principal component. Results did not significantly change depending on whether the post-stimulus time interval used for the analysis started 10 msec after the stimulus or at the peak or the end of the slow wave induced by the stimulation, nor did they change when the duration of the post-stimulus time interval increased from 10-800 msec to 10-1000 msec.

### Phase Locking Factor (PLF)

The instantaneous PLF was calculated as in (Palva et al., 2005; Pigorini et al., 2015) to determine how the stimulation affected the phase of ongoing oscillations across trials. To focus on phase coupling that could only be explained by the stimulus, we assumed a Rayleigh distribution of the PLF values during baseline (-600 to -100 ms), and then performed a statistical comparison with the baseline for each electrode. PLF values below threshold (α < 0.05) were set to zero. Initially, the spectral PLF contribution was calculated using a moving band pass filter window that ranged from 0 to 200 Hz (4 Hz width; 2 Hz superposition) and for each band the instantaneous PLF was calculated. In agreement with previous experiments in humans (Pigorini et al., 2015), this analysis revealed that the 8 to 40 Hz frequencies, encompassing the alpha and beta bands, but not the higher frequencies (40-200 Hz), were useful to distinguish between wake and NREM sleep, as well to distinguish between NREM sleep and REM sleep. Thus, all final PLF analyses used the 8 to 40 Hz frequency range.

### Detection of spontaneous and evoked slow waves

Detection of individual slow waves was performed as previously described (Vyazovskiy et al., 2007; Vyazovskiy et al., 2009a; Vyazovskiy et al., 2009b) on the spontaneous LFP signal during baseline sleep, recovery sleep after sleep deprivation (3PM to 5PM) and during the induction of slow waves by electrical stimulation. From the continuous recording, all NREM windows were extracted based on standard criteria for scoring vigilance states: wake was characterized by a low-voltage, high- frequency LFP activity and phasic muscle activity; NREM sleep was characterized by the occurrence of high-amplitude slow waves, spindles, and low tonic muscle activity; in REM sleep, cortical LFPs resembled those seen in wake but muscle tone was absent, with the exception of occasional twitches. Waveforms were detected using a bipolar transcortical arrangement (Vyazovskiy et al., 2009a) between deep and superficial LFPs cortical channels (layers 5-6 vs. layers 1-2). The signal was first filtered in the slow activity band (0.5-4 Hz; Chebyshev Type II filter) and all positive and negative peaks were detected. Slow waves were defined as positive deflections between two consecutive negative deflections below the zero-crossing with a duration of at least 100 ms, as in previous studies (Vyazovskiy et al., 2007). Only slow wave with an amplitude greater than the 75th percentile were used (Funk et al., 2017). Slow wave polarity reversal in the dorsoventral axis, along with the presence of a peak in high gamma power in mid-layer 5, and histology were used to estimate the electrode location (Chauvette et al., 2010; Funk et al., 2017; Senzai et al., 2019).

### Spike analysis

#### Preprocessing

Recordings were preprocessed separately with the CatGT command-line tool (github.com/billkarsh/SpikeGLX), performing 300–9000Hz band-pass filtering, global demultiplexing common average referencing and automatic artifact detection and removal with default parameters.

#### Spike sorting

For each animal, probe and each stimulation depth, the preprocessed recordings containing the wake, NREM sleep and REM sleep pulses were then concatenated into a single recording on which spike sorting was performed using the Kilosort2.5 algorithm (Pachitariu et al., 2016). Recordings for the sevoflurane condition were sorted separately. In order to account for fast and slow drift, the algorithm first performs a drift correction pre-processing step (Steinmetz et al., 2021): for each temporal batch, a fingerprint of the distribution of units along the probe is constructed from the histogram of spike amplitudes at each channel. This fingerprint is used to compute, for each temporal batch, the vertical offset of each channel relative to a template obtained from iterative averaging of the rigidly aligned fingerprints. The data used for sorting is then corrected using kriging interpolation. Since we did not observe significant drift on fast timescales, we used 8 sec batches (instead of the default 2 sec) to increase the reliability of the batches’ fingerprint. Besides the batch size, we used default values for all but two kilosort parameters: the projection thresholds *Th* were set to [12 10] instead of [10 5] and *lambda* was set to 50 instead of 10, as we observed that these values reduced the number of putative false positive spike detections in our data.

#### Postprocessing, curation and unit selection

We used Jennifer Colonnell’s fork of the Allen institute’s *ecephys_spike_sorting* toolbox to postprocess kilosort’s output (https://github.com/jenniferColonell/ecephys_spike_sorting). This allowed us to mark some of the clusters as noise based on their template’s spatial and temporal spread and remove the spikes occasionally double-counted by Kilosort. We then removed the remaining noise clusters using *phy* (https://github.com/cortex-lab/phy). Finally, we excluded all clusters with firing rate below 0.5 Hz. Overall, the total number of clusters throughout the probe selected for further analyses ranged from 65 to 402. Sorting data was extracted using the *SpikeInterface* toolbox (Buccino et al., 2020).

### OFF period detection

#### Peri stimulus time histograms (PSTH)

For the analysis of spikes locked to electrical or optogenetic stimulation, all time stamps corresponding to individual spike occurrences were concatenated across all recording channels showing single and/or multi-unit activity. 4 ms bin firing rate from -1 to +1 seconds, relative to stimulation time, was isolated and normalized to the baseline firing rate, defined as 1 to 0.4 seconds before the stimulation during wake.

#### Peri slow wave time histograms (PSWTH)

For the analysis of spikes locked to slow waves, all time stamps corresponding to individual spike occurrences were concatenated across all recording channels showing single and/or multi-unit activity. 4 ms bin firing rate from -1 to +1 seconds, relative to the slow-wave zero crossing, was isolated and normalized to the baseline firing rate, defined as 1 to 0.4 seconds before the slow wave zero crossing.

#### OFF periods

Using both PSTH and PSWTH, the time when the firing rate drops below 25% of the baseline was defined as the onset of the OFF period, while the start of the ON period was defined as the time when the firing rate rose above 25% of the baseline. OFF duration is equal to ON start time minus OFF start time.

### Current source density analysis (CSD)

For CSD analysis (Nicholson and Freeman, 1975; Chauvette et al., 2010) the following formula was applied:

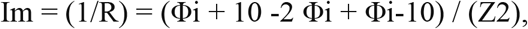

where Φi is the field potential in mV at a given electrode i, R is in MΩ, Z is the distance between electrodes in mm, Im is CSD in mV/mm2. Im > 0 and Im < 0 indicate Source (outward current) and Sink (inward current), respectively.

### Statistical analysis

The data were expressed as mean ± standard deviation. The significance of the differences among behavioral states was evaluated with repeated measures ANOVA, with the Greenhouse-Geisser correction, along with Tukey post hoc tests. For the comparison between waking and anesthesia a paired t-test was performed. A measure of effect size was reported for rmANOVA (η^2^) and t-test (d). The criterion used to reject null hypotheses was p < 0.05.

## Results

### Analysis of PCI^st^ in rats using electrical stimulation

PCI^st^ measurements started only after the sleep/waking pattern had normalized, usually at least one week after surgery. As expected since rats are nocturnal, animals spent most of the light period asleep and were mainly awake at night (Fig. 1A). Electrical stimuli were delivered only during the light period and trials occurred across several days to limit the number of stimuli delivered each day (Fig. 1A; ∼100 in each of the behavioral states: waking, NREM sleep, REM sleep). PCI^st^ was measured after electrical stimulation delivered by a laminar probe implanted perpendicular to the cortical surface, and recording was performed using one high-density Neuropixels probe (Fig. 1B). As in humans (Comolatti et al., 2019), PCI^st^ was derived from the averaged (across all trials) ERP for each of the cortical channels (∼ 80 to 100 channels in each rat), first by identifying the principal components that accounted for at least 99% of the response strength to the stimulation and then by calculating, for each component, the number of state transitions in the evoked response relative to the pre-stimulus baseline (Fig. 1C). Unlike in humans, however, all channels came from the same area (e.g. parietal association cortex, PtA). In almost all cases the electrodes spanned all layers of a given area, with the exception of the prefrontal cortex in which recordings came mainly from the deep layers.

**Figure 1.**
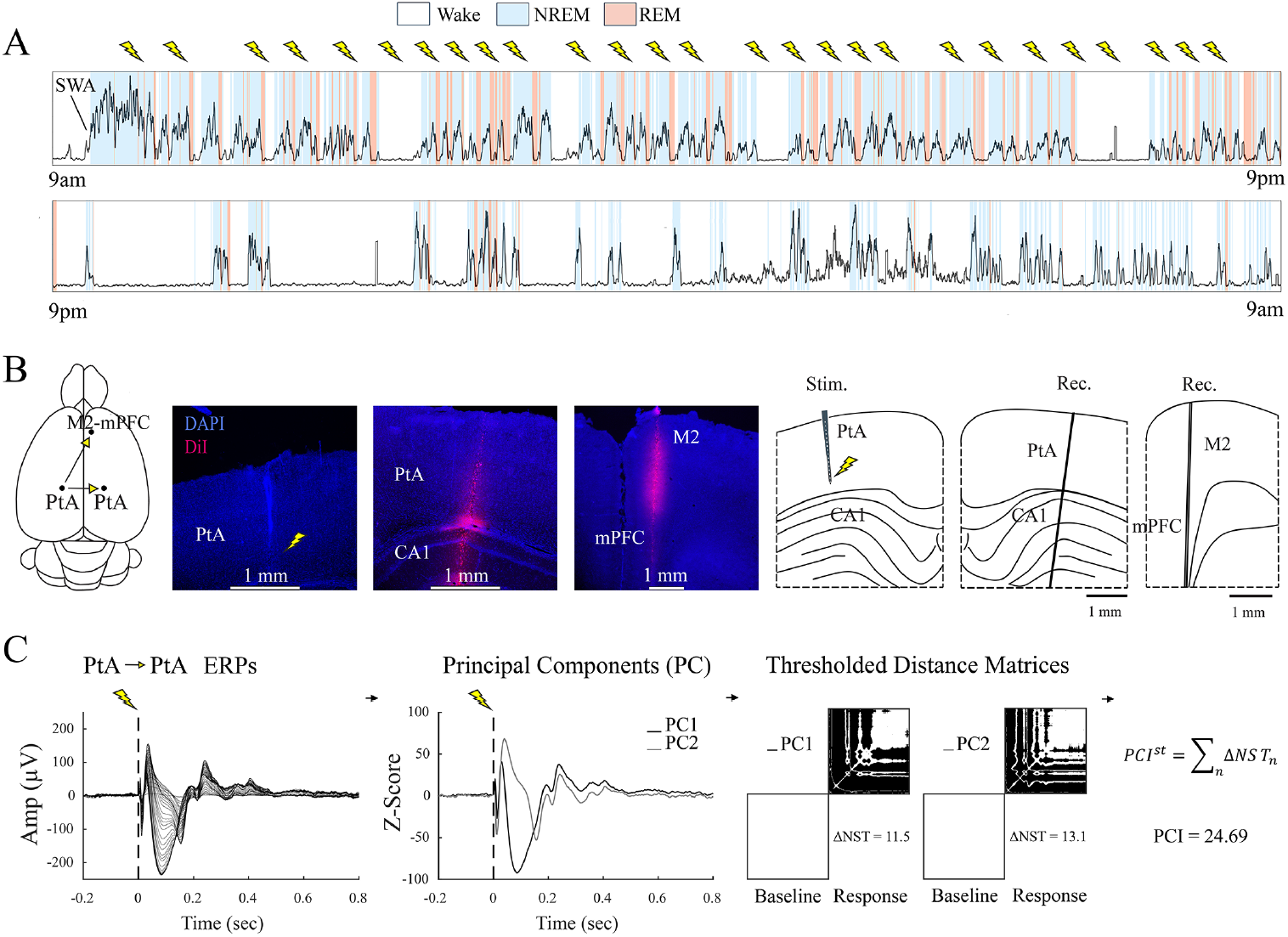
Experimental design. **A**, distribution of wake, NREM sleep and REM sleep during a continuous 24-hour recording in a representative rat. The black line shows slow wave activity (SWA, a. u.). As expected, SWA (the power in the 0.5 - 4 Hz range in cortical local field potentials) is elevated during NREM sleep and peaks at the beginning of the light period, the major sleep phase. Electrical stimuli (lightning bolts) were delivered during the light phase only. **B**, schematic of the rat brain displaying the position of the electrodes and coronal sections showing the location of stimulating and recording probes in one representative rat. **C**, left, averaged (across all trials) event related potentials (ERPs) for all cortical channels, each channel re-referenced to the white matter. To calculate PCI^st^, the ERPs for each cortical channel (81 channels in this example) are averaged across all trials (n=100 trials) and decomposed to identify the principal components (PC) of the ERPs. The “up and down” of each PC (state transitions, ST) are calculated after thresholding and compared between post- and pre- stimulation (distance matrix). PCI^st^ is the sum of the post/pre differences in ST (ΔNST) for all PCs (see Methods for details). In this and the following figures, the first (PtA in this example) and second (PtA in this example) cortical area indicate the site of stimulation and recording, respectively.

In the first set of experiments electrical stimuli were always delivered to the deep layers, and the cortical areas targeted for stimulation and recording varied across animals (Fig. 2A). In most cases, independent of the location of the stimulating and recording electrodes, rats (n = 8) showed more complex responses in wake and in REM sleep than in NREM sleep (Fig. 2B), leading to high PCI^st^ values in wake, low in NREM sleep, and intermediate or high in REM sleep in one or more of the recorded areas (Fig. 2A). In a few cases in which the recording electrode was contralateral and distant from the stimulating electrode (e.g. left M2 and right V2; left PtA and right M2) the evoked response was minimal or absent in one or more vigilance states, precluding PCI^st^ analysis or resulting in inconsistent changes across states (Fig. 2A). The number of principal components of the ERPs ranged from 1 to 4; most often there were 2 and the number did not change across behavioral states. Thus, changes in PCI^st^ were driven mostly by changes in the number of state transitions (Fig. 2B-C). PCI^st^ values in NREM sleep decreased on average by 50 ± 20% relative to wake and by 51 ± 27% compared to REM sleep, resulting in significant changes at the group level (paired ANOVA; F_(1.69, 23.7)_ = 33.4 ; p < 0.0001; ƞ^2^ = 0.70; Fig. 2D; Tukey correction for multiple comparison on Fig. 2D). The Phase Locking Factor (PLF), which measures for how long the evoked response remains phased locked to the original stimulus, was calculated for each recording site in wake and sleep. PLF was measured in the 8-40 Hz range because higher frequencies (40-200 Hz) did not discriminate across behavioral states (Fig. 2B), consistent with the results in humans (Pigorini et al., 2015). We found long PLF in wake and REM sleep and short PLF in NREM sleep (F_(1.80, 25.2)_ = 13.05 ; p = 0.0002; ƞ^2^ = 0.48) and PCI^st^ and PLF values were positively correlated (Fig. 2E). Grouping the recording sites in more anterior (M2, mPFC, Of) and more posterior (M1, PtA, V2) sites did not reveal any major difference, that is, significant sleep/wake differences in PCI^st^ could be detected in all areas (Fig. 2F; Frontal: F_(1.48, 10.4)_ = 14.84 ; p = 0.0015; ƞ^2^ = 0.67; Posterior: F_(1.38, 8.32)_ = 23.11 ; p = 0.0007; ƞ^2^ = 0.79). To assess the contribution of superficial and deep layers, for each recording site PCI^st^ was also calculated separately for the channels in the upper and lower half of the laminar probe. In general, both superficial and deep channels contributed to the sleep/wake changes (Fig. 2G; superficial: F_(1.58, 17.4)_ = 17.02 ; p = 0.0001; ƞ^2^ = 0.62; deep: F_(1.24, 13.6)_ = 6.47 ; p = 0.019; ƞ^2^ = 0.37).

**Figure 2.**
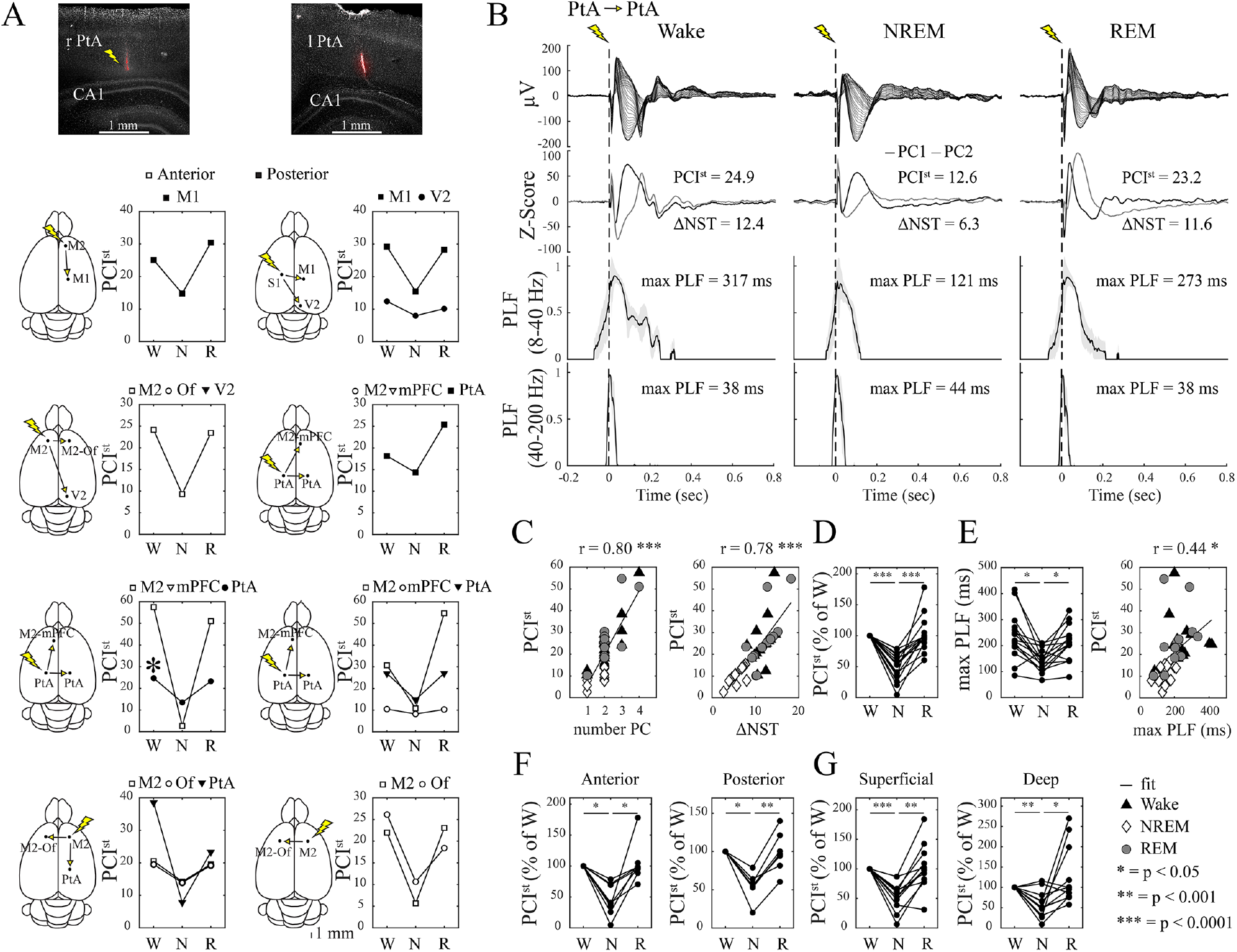
Sleep/wake changes in PCI^st^ in rat cortex. **A**, histology in a representative rat and schematic location of stimulating and recording electrodes in each animal, with corresponding PCIst values in wake (W), NREM sleep (N) and REM sleep (R). Empty and filled symbols indicate more anterior and more posterior regions, respectively, where PCI^st^ was measured. Cases in which ERPs were absent in wake are not included. M1, primary motor; M2, secondary motor; mPFC, prefrontal; Of, orbitofrontal; PtA, parietal association; V2, secondary visual. **B**, example of ERPs, their principal components (PC), and phase locking factor (PLF) for one rat (PtA, * in panel A). PLF is shown separately for the 8-40 Hz range and the 40-200 Hz range. The latter was not used in the main analysis because it does not discriminate across behavioral states. **C**, number of PC (left) and changes in the number state transitions ((ΔNST, right) for all experiments. Note that in most experiments PC = 2, independent of sleep and wake. **D**, group level changes in PCI^st^ across waking and sleep. **E**, group level changes in max PLF across waking and sleep and correlation with PCI^st^. **F**,**G** group level changes in PCI^st^ across wake and sleep shown separately for recording in anterior and posterior cortical regions, and for superficial and deep channels.

In 8 rats PCI^st^ was also compared between wake and deep anesthesia, with loss of righting reflex, induced using sevoflurane (2%; 13 areas) or dexmedetomidine (100ug/kg; 6 areas) (Fig. 3A). In all cases, independent of the location of stimulating and recording electrodes, PCI^st^ was lower under anesthesia compared to waking and the difference was mainly driven by changes in the number of state transitions (Fig. 3B,C). In most cases PLF values were lower under anesthesia and were positively correlated with PCI^st^ values (Fig. 3D). PCI^st^ values in anesthesia did not differ significantly from those during NREM sleep (W, NREM, A; F _(1.74, 27.9)_ = 66.7; p < 0.0001; ƞ^2^ = 0.81; NREM vs A, p = 0.1267).

**Figure 3.**
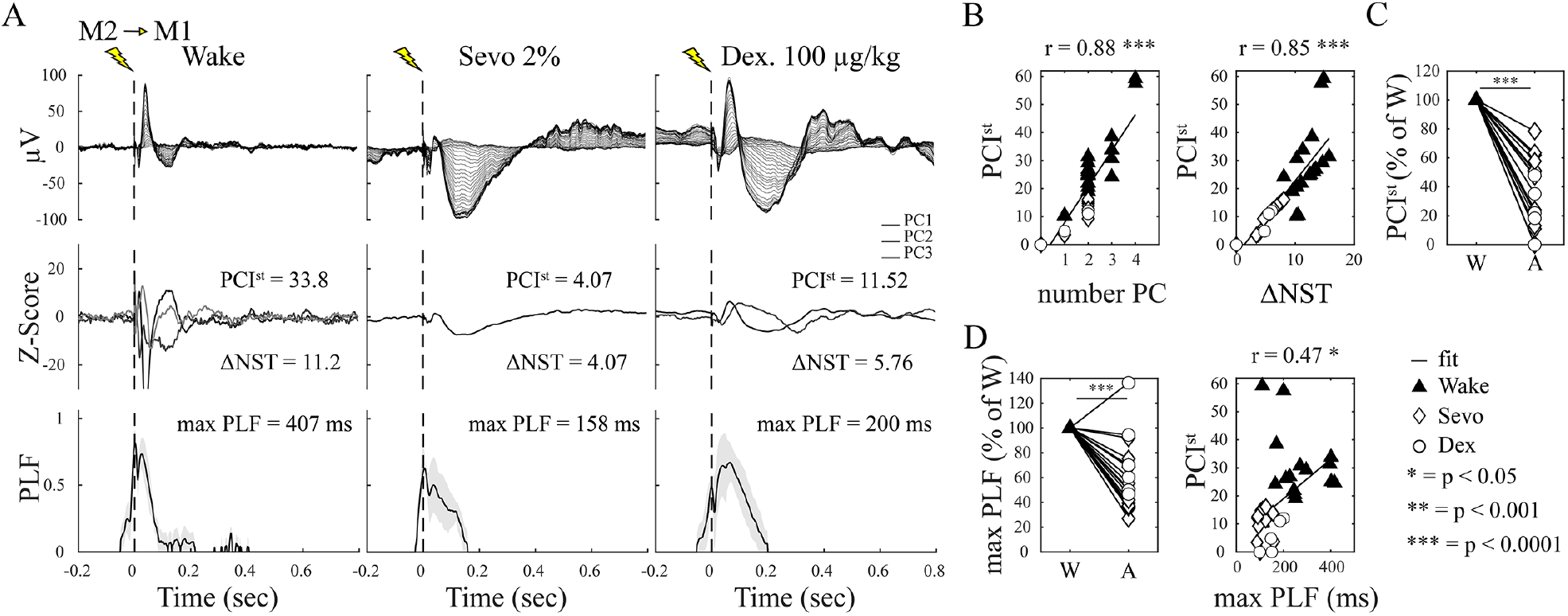
Anesthesia-induced changes in PCI^st^ in rat cortex. **A**, example of ERPs (top), their principal components (PC, middle), and phase locking factor (PLF, bottom) for one representative rat. **B**, number of PC (left) and changes in the number of state transitions ((ΔNST, right) for all experiments. **C**, group level changes in PCI^st^ between wake and anesthesia. Cases in which ERPs were absent in wake are not included. **D**, group level changes in max PLF between wake and anesthesia and correlation with PCI^st^.

In 4 rats the stimulation was delivered at different depths spanning superficial and deep layers, and the response was measured across all layers (Fig. 4A,B). ERPs in both ipsilateral and contralateral cortex were large when deep layers were stimulated and small or undetectable when the stimulation was restricted to the most superficial layers (Fig. 4C,D). As a result, in both ipsilateral and contralateral cortex the sleep/wake changes in PCI^st^ were robust for stimulation of deep and middle layers, but inconsistent when only the most superficial layers were stimulated (Fig. 4E,F).

**Figure 4.**
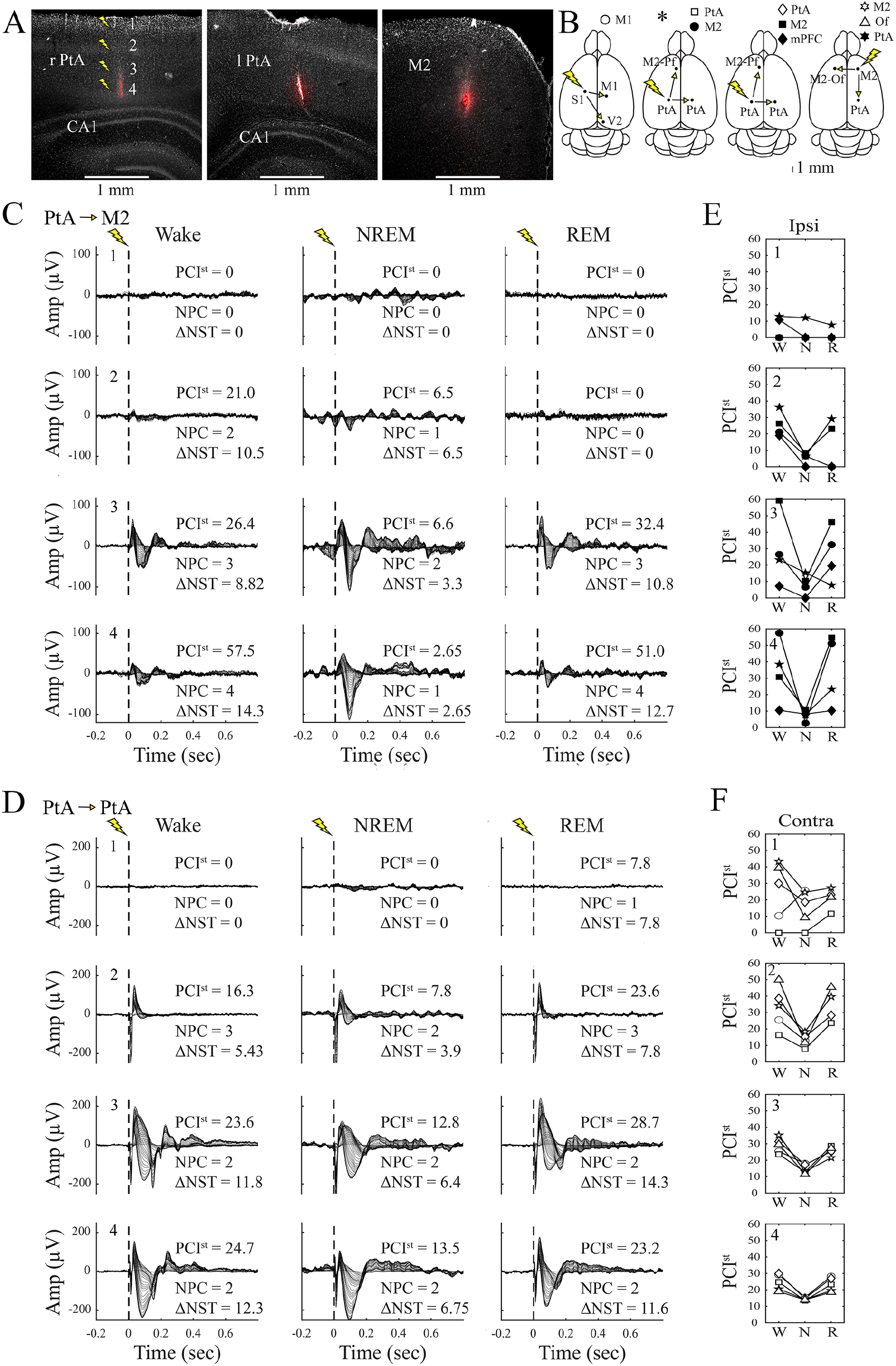
Changes in PCI^st^ depending on the depth of stimulation. **A**, histology in a representative rat. **B**, schematic location of stimulating and recording electrodes in the 4 rats used for depth analysis. **C**, example of ERPs and PCI^st^ values after stimulation in PtA at 4 different depths and recording in ipsilateral M2 in one rat (asterisk in panel B). **D**, example of ERPs and PCI^st^ values after stimulation in PtA at 4 different depths and recording in contralateral PtA in one rat (asterisk in panel B). **E-F**, sleep/wake changes in PCI^st^ for all 4 rats, shown separately depending on depth of stimulation. The different symbols refer to the areas indicated in panel B, with filled and open symbols indicating ipsilateral and contralateral stimulation, respectively.

To test whether electrical stimulation triggers an OFF period, Neuropixels recordings were spike- sorted using the Kilosort2.5 algorithm (Pachitariu et al., 2016), followed by manual curation (see Methods). We focused on 6 rats that had a good yield of cortical units (67 ± 42 single units, 38 ± 31 MUA per cortical area, mean ± std dev) and in which electrical stimulation produced robust ERPs in all 3 states, resulting in reliable changes in PCI^st^. For each rat and each area separately, we first analyzed sleep in baseline and after 6 hours of sleep deprivation, without electrical stimulation, to define the range in the duration of the OFF periods. From the peri-slow-wave time histograms corresponding to all areas (Fig. 5A) we obtained an average duration of OFF periods of 48 ± 14 ms (mean ± std dev; range 35-68 ms) during baseline sleep and of 97 ± 37 ms (range 48-152 ms) during the first 2 hours of recovery sleep after sleep deprivation. We then tested whether the electrical stimulation triggered a bona fide OFF period, as defined based on the analysis in sleep, and calculated its duration for each experimental condition (wake, NREM sleep, REM sleep). In 5 out of 6 rats periods of neuronal silence (62 ± 27 msec, mean ± std dev) were induced during NREM sleep (Fig. 5A) and their amplitude and laminar distribution were very similar to those of the spontaneous OFF periods (Fig. 5B). This pattern applied to all areas examined (M1, M2, PtA). OFF periods were either absent in wake and REM sleep or, if present, were short lasting. Moreover, these occasional OFF periods were followed by a rebound in firing that exceeded the pre-stimulation levels of spiking, which was not the case for the OFF periods evoked in NREM sleep. In two rats in which OFF periods were present in all three behavioral states (albeit longer in NREM sleep), the rebound firing was still present only in wake and REM sleep (Fig. 5A). In a few cases no clear OFF periods in NREM sleep were observed, but the mean firing rate still showed a decline after the electrical stimulation, while no clear decrease in unit activity occurred in wake and REM sleep. In summary, OFF periods occur in most cases after stimulation during NREM sleep. OFF periods can happen in wake and REM sleep, but they are shorter and, unlike those in NREM sleep, they are usually followed by a strong rebound in unit firing. Finally, in a few animals with high yield of both cortical and thalamic units the single-unit peri- stimulus time histogram showed that electrical stimulation during NREM sleep triggered a period of strongly reduced activity in both cortex and thalamus. Notably, however, the thalamic OFF period was shorter, leading to an earlier rebound of firing in thalamus than in cortex, a pattern also seen during physiological NREM sleep (Fig. 5C). In wake and REM sleep the thalamic OFF period was shorter than in NREM sleep and was not followed by a clear rebound in firing.

**Figure 5.**
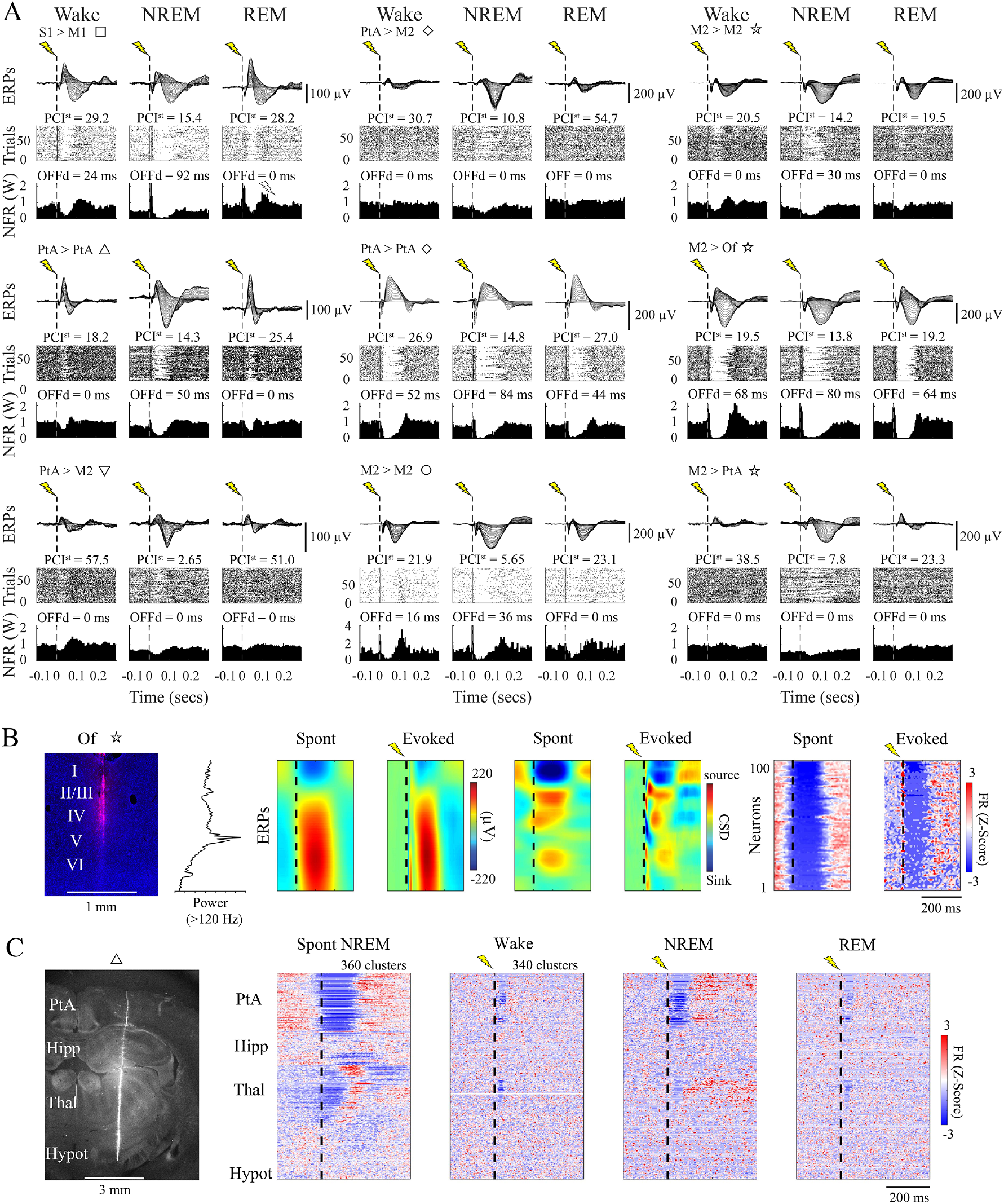
OFF periods triggered by electrical stimulation in rats. **A**, Nine examples of ERPs in 6 rats (top), corresponding changes in firing rate across all trials (middle) and mean changes in firing rate (normalized to the wake values, NFR). In each example, the first and second cortical area indicate the site of stimulation and recording, respectively, followed by a symbol that identifies the specific animal. OFFd, duration of the OFF period. **B**, comparison between spontaneous and evoked slow waves in orbitofrontal cortex (Of) showing similar amplitude, laminar distribution (CSD, current source density) and unit activity (FR, firing rate). Here and in C, time zero (vertical dashed line) represents slow-wave zero crossing (spont) and electrical stimulation (evoked). Histology and the peak in gamma (>120Hz) power in mid-layer 5 were used to estimate the electrode location (left panels). **C**, example of single-unit peri-stimulus time histograms (PSTH) locked to the slow wave zero-crossing (spontaneous NREM sleep) or the electrical stimulation in wake, NREM sleep and REM sleep (505 clusters; 80 pulses) in a parietal probe (PtA), sorted by depth. Each row corresponds to the PSTH of a single unit. For each cluster, evoked rates were zscore-normalized based on the mean and standard deviation of instantaneous evoked rates across bins from -2sec to -10msec before the pulses. Region boundaries were obtained from histological reconstruction (left panel); Cx, cortex; Hipp, hippocampus; Th, thalamus; Hypot, hypothalamus.

### Analysis of PCI^st^ in mice using optogenetic stimulation

Cortical stimuli were delivered at least 1-2 weeks after surgery to allow the sleep/wake pattern to normalize. Mice, like rats, were asleep mainly during the day and had several hours of spontaneous wake at night, and stimuli were delivered only during the light phase (Fig. 6A). In CaMKIIα::ChR2 mice the excitatory opsin is expressed in the pyramidal neurons of the cortex and hippocampus (Stark et al., Neuron 2014). For the stimulation of the posterior parietal association area (PtA), the optic fibers were placed on the cortical surface to avoid the stimulation of the hippocampus. For anterior stimulation, optic fibers were implanted deep in prefrontal cortex to target as much as possible all layers (Fig 6B). ERPs recorded from PtA and other areas outside the prefrontal cortex (V2, S1) showed a pattern consistent with the one seen in rats: responses were complex in wake and REM sleep and tended to be larger but more stereotyped in NREM sleep (Fig. 6C). This pattern occurred independent of the stimulated area (PtA or prefrontal cortex) and resulted in PCI^st^ values that were higher in wake than in NREM sleep, and almost always higher in REM sleep than in NREM sleep (Fig. 6D,G; F_(1.35, 8.14)_ = 6.23 ; p = 0.0302; ƞ^2^ = 0.51). By contrast, ERPs recorded from frontal and prefrontal cortex were either small or stereotyped across states (Fig. 6E), and led to PCI^st^ values that did not differ significantly across states but were higher in NREM sleep than in wake (Fig. 6D,G; F_(1.08, 3.24)_ = 2.21 ; p = 0.2298; ƞ^2^ = 0.42).

**Figure 6.**
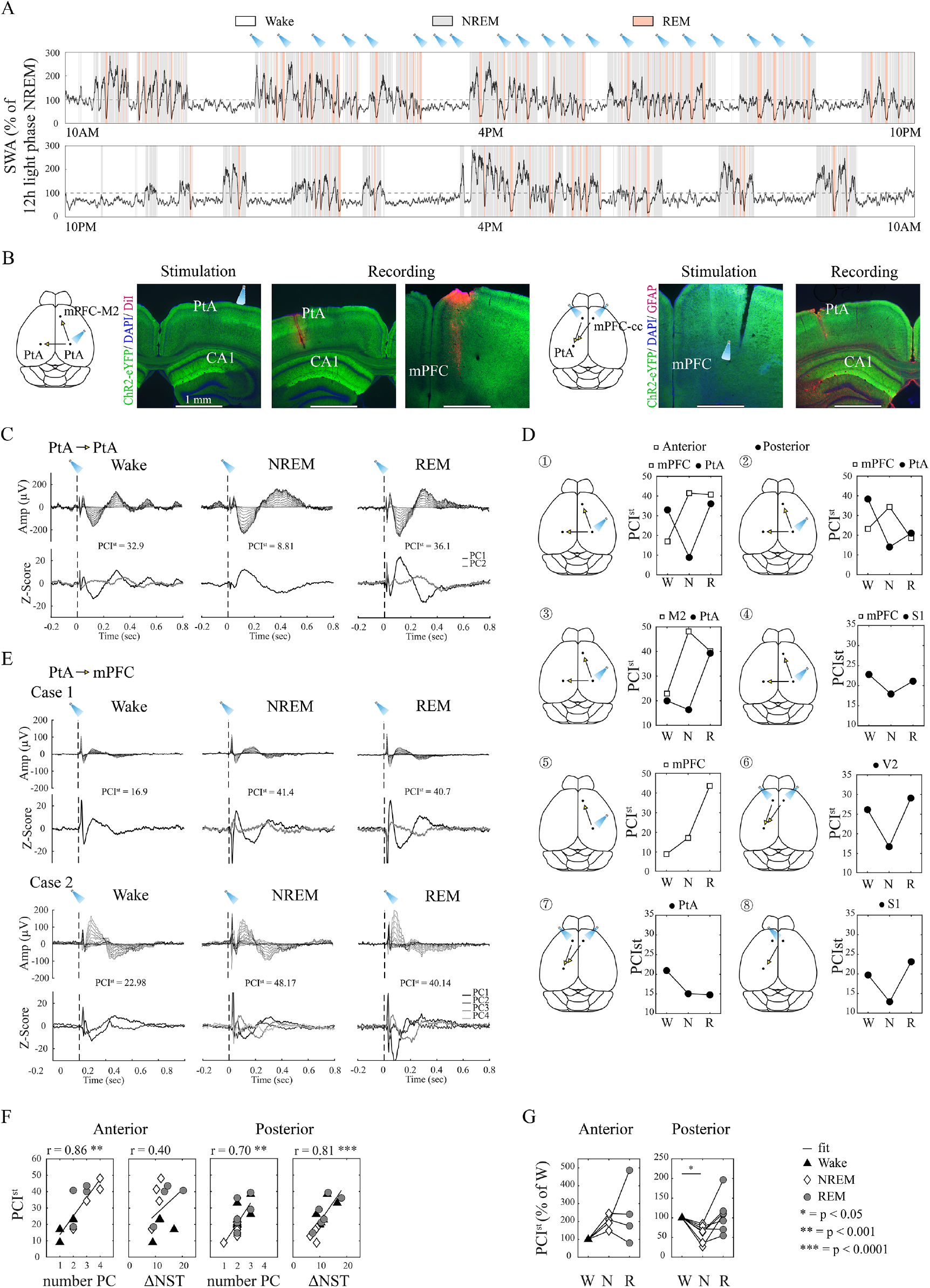
Sleep/wake changes in PCI^st^ in mouse cortex. **A**, distribution of wake, NREM sleep and REM sleep during a continuous 24-hour recording in a representative mouse. The black line shows relative slow wave activity (SWA), which as in rats peaks at the beginning of the light period, Optogenetic stimuli were delivered during the light phase only. **B**, schematic of the mouse brain displaying the position of the electrodes and coronal sections showing the location of stimulating and recording sites in two mice with stimulation in frontal or posterior cortex. DAPI and GFAP (glial fibrillary acidic protein) staining were used to identify cortical layers and probes, respectively. **C**,**E**, examples of ERPs (top) and their principal components (PCs, bottom) for 3 mice. **D**, schematic location of stimulating and recording electrodes in each animal, with corresponding PCIst values in wake (W), NREM sleep (N) and REM sleep (R). Empty and filled symbols indicate more anterior and more posterior regions, respectively, where the recording electrode was located. Cases in which ERPs were absent in wake are not included. S1, primary sensory; M2, secondary motor; mPFC, medial prefrontal; PtA, parietal association; V2, secondary visual. **F**, number of PCs and changes in the number of state transitions for all experiments, shown separately for anterior and posterior regions. **G**, group level sleep/wake changes in PCI^st^ recorded in anterior and posterior regions.

In 9 mice PCI^st^ was also measured in deep anesthesia (sevoflurane, 1-2%; dexmedetomidine 70-100μg/kg) (Fig. 7A-C). ERPs recorded from PtA and other posterior areas showed complex responses in wake and more stereotyped responses in anesthesia, while ERPs recorded from frontal and prefrontal cortex were often small or stereotyped in all cases (Fig. 7A,B). PCI^st^ values recorded in posterior areas were always lower in anesthesia than in wake, while in anterior areas they were inconsistent (Fig. 7D).

**Figure 7.**
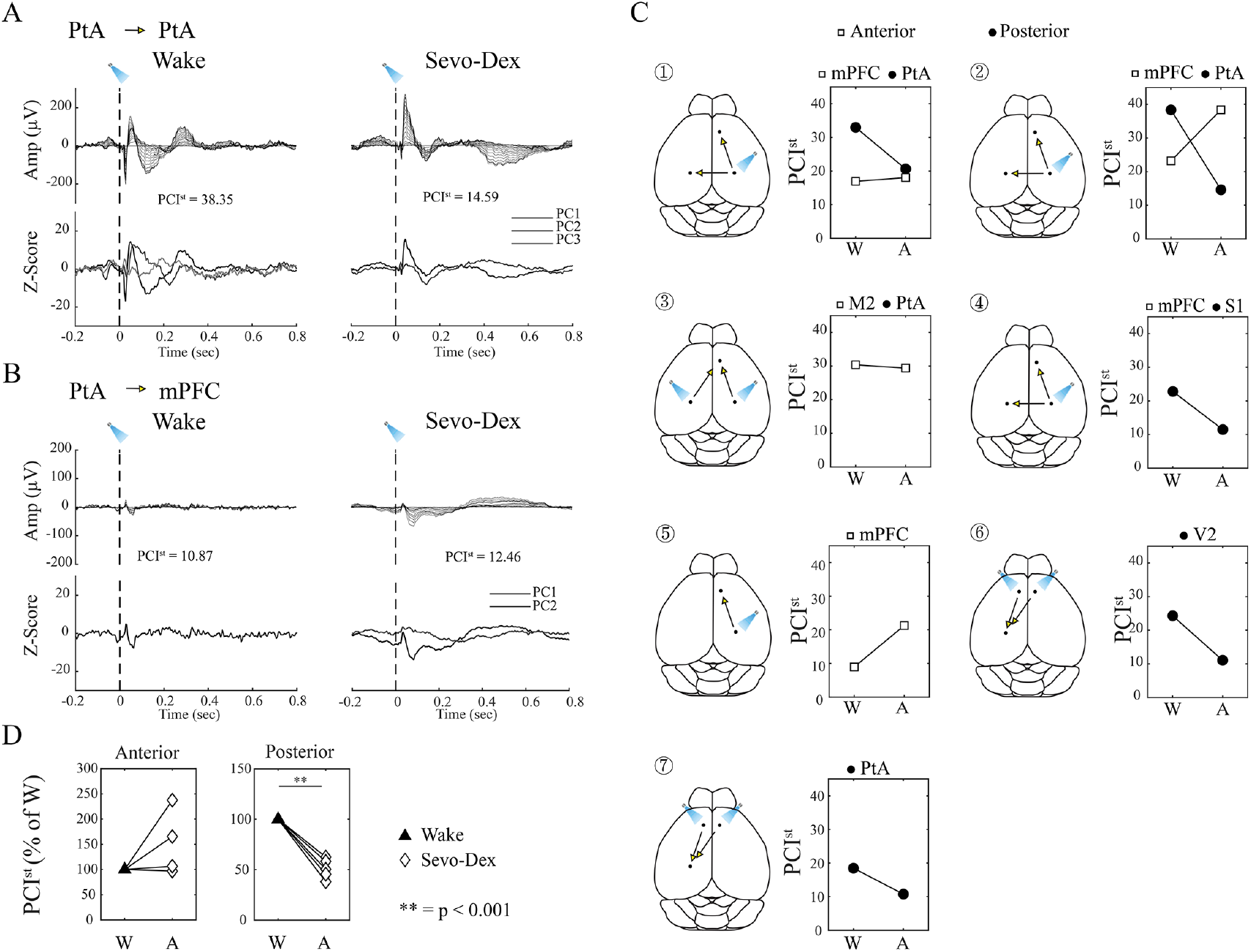
Anesthesia-induced changes in PCI^st^ in mouse cortex. **A,B** example of ERPs and their principal components (PC) for two mice. **C**, schematic location of stimulating and recording electrodes in each animal, with corresponding PCIst values in wake (W) and anesthesia. Empty and filled symbols indicate more anterior and more posterior regions, respectively, where the recording electrode was located. Mice are arranged following the order in Figure 6 (anesthesia data are missing in 2 mice). Areas are labelled as in Figure 6. Cases in which ERPs were absent in wake are not included. **D**, group level changes in PCI^st^ between wake and anesthesia.

**Figure 8.**
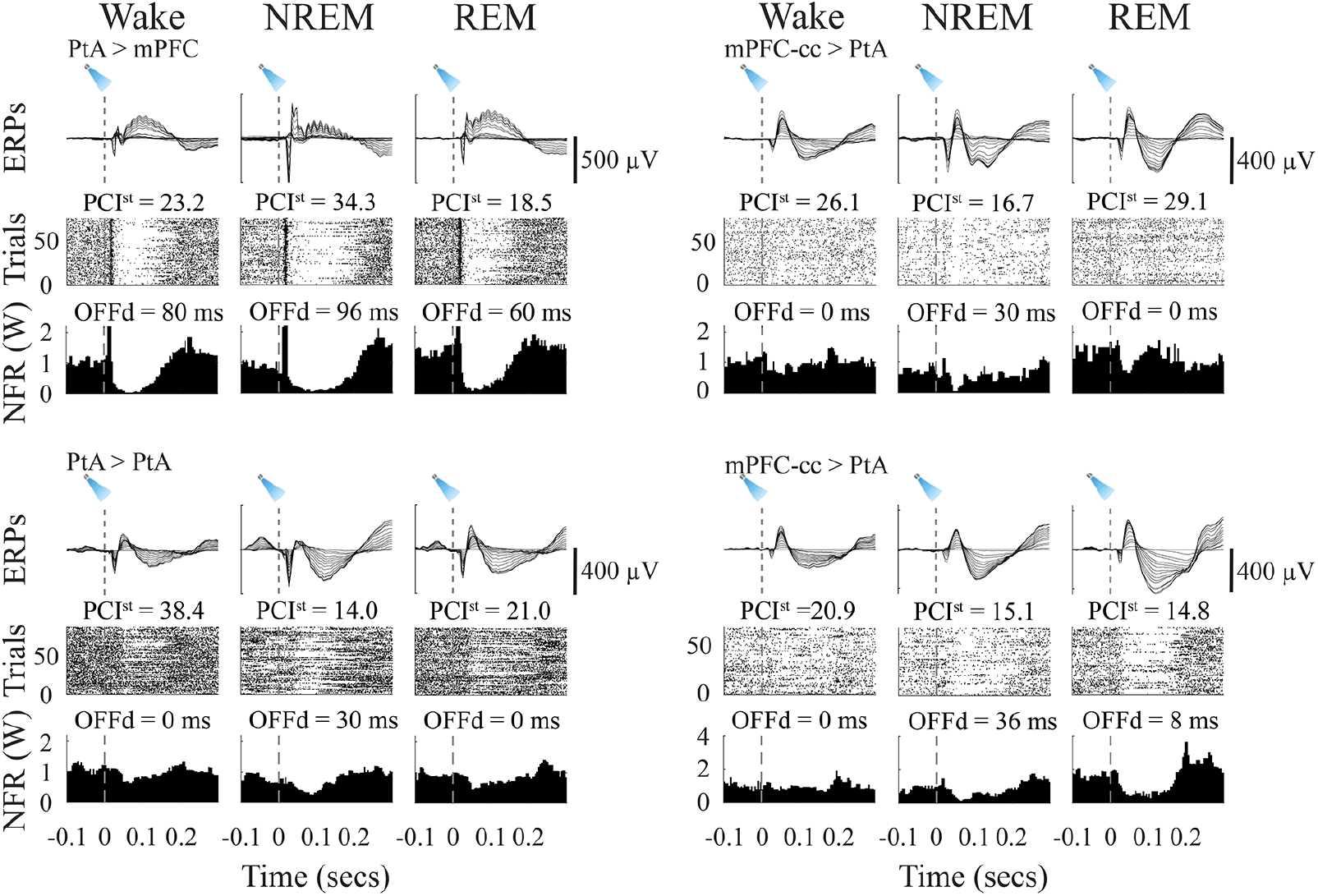
OFF periods triggered by optogenetic stimulation in mice. Four examples of ERPs (top), corresponding changes in firing rate across all trials (middle) and mean changes in firing rate (normalized to the wake values, NFR). In each example, the first and second cortical area indicate the site of stimulation and recording, respectively. OFFd, duration of the OFF period.

Finally, we tested whether the optogenetic stimulation triggers OFF periods also in mice. Independent of whether the stimulation was anterior or posterior, OFF periods were consistently recorded from PtA during NREM sleep but were absent in wake; during REM sleep OFF periods were either absent or, when present, they were followed by a rebound firing that exceeded the pre-stimulation levels. In prefrontal cortex, by contrast, long OFF periods followed by strong rebound firing occurred in all states.

## Discussion

Measures of perturbational complexity, such as PCI and PCI^st^, can be used to assess the presence and absence of consciousness without relying on behavioral reports. These indices have been validated in a large number of human subjects in many different conditions (Casarotto et al., 2016) and have shown unrivaled sensitivity and specificity (Sarasso et al., 2021). They have also proven their value in inferring the presence and absence of consciousness in unresponsive patients (Casarotto et al., 2016).

Our first goal was to validate the use of PCI^st^ in animal models of spontaneous sleep and wake. We found that PCI^st^ is high in wake and REM sleep and low in NREM sleep in both rats and mice, as it is in humans (Casali et al., 2013; Casarotto et al., 2016). In humans, wakefulness is invariably conscious, and REM sleep is most often accompanied by dreaming. By contrast, consciousness frequently fades during NREM sleep, especially early in the night, when the EEG shows high amplitude slow waves, especially in posterior cortex (Siclari et al., 2017). The similarity of the results obtained in freely moving rodents suggests that PCI^st^ may be used as a reliable readout of the effectiveness of causal interactions in corticothalamic networks that are thought to underlie the capacity for experience (Tononi et al., 2016).

We also found that, as in humans, PCI^st^ is reduced in rats and mice under deep slow wave anesthesia with sevoflurane and/or dexmedetomidine. These results confirm and extend the findings of a recent study in head fixed rats (Arena et al., 2021). In that study, propofol and sevoflurane anesthesia induced large slow waves and led to a decrease in PCI^st^ associated with decreased phase-locking, whereas ketamine anesthesia was associated with wake-like EEG activity and with PCI^st^ values intermediate between wake and propofol/sevoflurane. Thus, in both humans and rodents, conditions characterized by the presence of widespread cortical slow waves (deep NREM sleep, propofol, sevoflurane and dexmedetomidine anesthesia) are associated with low PCI^st^ values, while wake-like EEG activity is associated with intermediate or high PCI^st^ (ketamine anesthesia, REM sleep, wake).

Which cellular and network mechanisms underly the ability of measures of perturbational complexity to reflect the level of consciousness? Theoretical considerations predict that the loss of consciousness should be associated with a breakdown of causal interactions within corticothalamic networks (Tononi et al., 2016). In patients, deep intracranial stimulation during wake triggered complex and long-lasting cortical evoked responses that were deterministically linked to the initial stimulus (Pigorini et al., 2015). By contrast, the same stimulation during NREM sleep (REM sleep was not studied) triggered a suppression of high frequencies (>20Hz) and an associated increase in low frequencies (<4Hz). Moreover, when cortical activity resumed, it was not phased-locked to the original stimulus, and the cessation of phase locking was correlated in time with the suppression of high frequencies (Pigorini et al., 2015). However, due to the unavailability of unit recordings in humans (Pigorini et al., 2015) and rats (Arena et al., 2021), it could not be determined whether the drop in high frequencies and the loss of a deterministic response during NREM sleep reflects the occurrence of a period of neuronal silence due to neuronal bistability.

The present recordings in freely moving rats show that phase locking to electrical stimulation was also long in wakefulness and REM sleep and short in NREM sleep. Moreover, unit recordings in both rats and mice demonstrate that low PCI^st^ values during NREM sleep and slow wave anesthesia are indeed associated with the early occurrence of OFF periods. This provides direct support to the hypothesis that sustained causal interactions that lead to high PCI^st^ cannot take place when cortical networks are bistable. In some cases, electrical or optogenetic stimulation during wake or REM sleep also triggered neuronal silence, but the OFF period was shorter than in NREM sleep. Intriguingly, these OFF periods were followed by a strong rebound in cortical firing that was absent in NREM sleep. A local, low-amplitude, short-lasting increase in low frequencies (< 4Hz) after deep intracranial stimulation can also occur during wakefulness in humans, while the suppression of high frequencies only occurs during NREM sleep (Pigorini et al., 2015). Previous evidence indicates that OFF periods may be triggered by Martinotti cells that powerfully inhibit every other neuronal population (Funk et al., 2017). Thus, strong local stimulation may activate Martinotti cells strongly enough to trigger local OFF periods even in wakefulness (Funk et al., 2017).

In humans, phase locking was suppressed during NREM sleep predominantly in the alpha and beta bands (8-30 Hz), while phase locking in the 30-100 Hz gamma band was short-lasting and comparable in wake and NREM sleep (Pigorini et al., 2015). In our study, phase locking was suppressed in NREM sleep in the 8-40 Hz band but not in the gamma band (40-200 Hz), compared to both wake and REM sleep. Alpha and gamma oscillations have been linked to feedback and feedforward oscillations, respectively (van Kerkoerle et al., 2014). Since PCI^st^ and phase locking values are positively correlated, it may be that the longer phase locking during wake and REM sleep relative to NREM sleep is associated with the activation of feedback loops in cortico-cortical circuits and may sustain the higher spatiotemporal complexity observed in conscious states (Pigorini et al., 2015).

In humans, PCI^st^ values are calculated based on EEG or intracranial signals coming from multiple cortical sites. The recent study in anesthetized rats also calculated PCI^st^ using a grid of 16 EEG screws covering bilaterally most of the dorsal cortex (Arena et al., 2021). Here, we found that changes in perturbational complexity could be reliably estimated from a single recording probe, albeit one endowed with multiple contacts across its length. We hypothesize that high PCI^st^ values during wake and REM sleep may reflect the triggering of complex reverberatory activity across multiple cortico- thalamic and cortico-cortical loops that impinge on different contacts at different times. By contrast, during NREM sleep, and even more so during anesthesia, this reverberatory activity may be blocked by the widespread occurrence of OFF periods. As shown in Fig. 5B, it is sometimes possible to document the triggering of reverberatory activity, in this case a cortico-thalamic volley followed by a cortico-thalamic OFF period, which is brief in wake and REM sleep. This is followed by rebound spiking occurring first in thalamic neurons, possibly triggering secondary cortical activity that is complex and long-lasting in wake and REM sleep, but localized and short-lasting in NREM sleep (see also (Crunelli and Hughes, 2010; Urbain et al., 2019)).

In rats, the stimulation of both anterior and posterior cortex provided consistent PCI^st^ results when applied to the deep and middle layers but not when delivered to the most superficial layers. Large, thick- tufted pyramidal cells of L5b are mainly involved in cortico-subcortical loops, and layer 6 pyramidal cells project to the thalamus, while pyramidal cells in L2/3 have the densest cortico-cortical anatomical connections compared to infragranular layers (Harris and Shepherd, 2015). Thus, the stimulation of neurons in the deep and middle layers is more likely to recruit cortico-thalamic and other cortico- subcortical loops, increasing the probability that a single distant recording site detects a complex evoked response during wake.

On the other hand, at the recording site both superficial and deep channels contributed to high PCI^st^ in wake and REM sleep and low PCI^st^ in NREM sleep and anesthesia. It is currently unknown whether supragranular or infragranular layers, all cortical layers, or only specific cellular populations are especially important to account for the presence and content of consciousness. In humans, a negative slow cortical potential likely originating from supragranular layers (He and Raichle, 2009) appears between stimulus onset and behavioral response only when a near-threshold stimulus is perceived (Pins and Ffytche, 2003) and has been proposed as ‘generalized awareness negativity’, a physiological correlate of consciousness across sensory domains (Dembski et al., 2021). In mice, on the other hand, deep general anesthesia decouples the signaling from the apical dendrites to the cell body of layer 5 but not of layer 2/3 pyramidal neurons (Suzuki and Larkum, 2020). Thus, loss of consciousness under anesthesia was associated with the impaired activity of pyramidal neurons in deep but not in superficial layers.

An intriguing observation is that in rats PCI^st^ changed across vigilance states (high in wake and REM sleep, low in NREM sleep and anesthesia), regardless of the site of stimulation and of whether the recording electrode was placed in anterior or posterior cortex. In mice results were similar, except when recording from prefrontal electrodes, which showed inconsistent PCI^st^ changes with behavioral state. This may be because neural circuits in frontal/prefrontal areas are less developed in mice than in rats (Van De Werd et al., 2010). However, a key methodological difference is that in rats we used high-intensity electrical stimulation (as in humans), likely recruiting a broad cortical network and fibers of passage. In mice, we used instead optogenetic stimulation to selectively target neighboring excitatory pyramidal neurons. This was adequate to trigger complex responses from posterior cortex during wake and REM sleep, but not from anterior cortex. Perhaps anterior areas may be organized in a way less suitable for sustaining causal interactions that are both integrated and differentiated, and thereby consciousness, in line with lesion, stimulation, and recording studies (Boly et al., 2017).

Overall, our experiments show that measures of perturbational complexity can be used for the reliable assessment of vigilance state in rodents. In humans, purely behavioral readouts, even refined ones such as the Glasgow Coma scale revised, administered by expert neurologists, result in a substantial proportion of false negatives in unresponsive patients. In animals, behavioral readouts such as the righting reflex are even more difficult to evaluate (Fuller et al., 2011; Gao and Calderon, 2020). Thus, PCI^st^ may offer a promising proxy for assessing consciousness in animals, with potential benefits in terms of research ethics and well-being.

## Acknowledgements and Funding sources

Supported by U.S. Department of Defense grant W911NF1910280 (CC, GT), NIH grant 1R01GM116916 (GT), the Tiny Blue Dot Foundation (GT) and the Templeton World Charity Foundation (CC).

## Notes

**Conflict of interest statement:** The authors declare no competing financial or non-financial interests.

### Competing Interest Statement

The authors have declared no competing interest.

